# rRNA transcription is integral to liquid-liquid phase separation and maintenance of nucleolar structure

**DOI:** 10.1101/2022.11.14.516489

**Authors:** Soma Dash, Maureen C. Lamb, Jeffrey J. Lange, Mary C. McKinney, Dai Tsuchiya, Fengli Guo, Xia Zhao, Timothy J. Corbin, MaryEllen Kirkman, Kym Delventhal, Emma L. Moore, Sean McKinney, Rita Shiang, Paul A. Trainor

## Abstract

Beginning with transcription of ribosomal RNA (rRNA) by RNA Polymerase (Pol) I in the nucleolus, ribosome biogenesis is intimately tied to cell growth and proliferation. Perturbation of ribosome biogenesis has been previously shown to affect nucleolar structure, yet the underlying mechanism is unknown. We generated loss-of-function mouse mutants of Pol I subunits, *Polr1a, Polr1b, Polr1c* and *Polr1d*, thereby genetically inhibiting rRNA transcription and ribosome biogenesis. Pol I mutant embryos are preimplantation lethal and have fewer nucleoli. Using hiPSCs triple labeled for the three nucleolar compartments, we observe two phenotypes upon Pol I inhibition: a single condensed nucleolus, and fragmented nucleoli. We find that when rRNA transcription is inhibited, the viscosity of the granular compartment of the nucleolus is increased disrupting its liquid-liquid phase separation properties, which results in a condensed nucleolus. Taken together, our data suggests that Pol I function and rRNA transcription are required for maintaining nucleolar structure and integrity.

## Introduction

Ribosome biogenesis is a fundamental process required for protein synthesis in all cells, and therefore, is vital for all cell survival and proliferation^1^. Ribosome biogenesis takes place in the nucleolus, which is a membrane-less organelle formed around actively transcribing ribosomal DNA (rDNA) loci in the nucleus of a cell^2–6^. The nucleolus is tripartite in structure with each compartment having a specific function in the process of ribosome biogenesis^7–9^. The fibrillar center (FC) constitutes the innermost compartment of the nucleolus and forms around active rDNA. It is surrounded by the dense fibrillar center (DFC), and rDNA is transcribed into pre-ribosomal RNA (rRNA) by RNA Polymerase (Pol) I together with associated proteins such as UBF, Treacle and Nucleolin (NCL) at the boundary of the FC and DFC. Pre-rRNA is then transported into the DFC where it is cleaved and modified by rRNA processing proteins such as Fibrillarin (FBL) and Nopp140. The granular component (GC) surrounds the FC and DFC and is where the assembly of processed rRNA with ribosomal proteins (RP) occurs, the process of which is mediated by proteins such as Nucleophosmin1 (NPM1)^10,11^. The different compartments of the nucleolus are maintained through liquid-liquid phase separation (LLPS), in which immiscible liquid phases sort into distinct and defined regions^8^. LLPS relies on weak interactions between RNA and proteins with intrinsically disordered regions (IRDs)^12,13^, such as FBL^14^ and NPM1^15^, to concentrate particular proteins and RNA to a specific compartment of the nucleolus. Importantly, previous studies demonstrate that changes in the LLPS properties of a nucleolar compartment alters the morphology and function of the nucleolus^16^.

In the nucleolus, Pol I is responsible for rRNA transcription which is a critical rate-limiting step in the ribosome biogenesis process^17,18^. Pol I consists of thirteen subunits, of which Polr1a and Polr1b form the catalytic core, while Polr1c and Polr1d form a clamp holding the catalytic core together^19^. Mutations in Pol I subunits cause developmental defects that disproportionally affect craniofacial development. More specifically, mutations in *POLR1A* cause Acrofacial Dysostosis Cincinnati type (AFDCin) while mutations in *POLR1B, POLR1C* and *POLR1D* and the Pol I associated protein, *TCOF1*/TREACLE lead to Treacher Collins Syndrome (TCS)^20–24^. We have previously identified the mechanisms underlying the pathogenesis of craniofacial defects in association with Pol I loss-of-function. In Pol I subunit mutants, rRNA transcription is impaired which alters the stoichiometry between rRNAs and RPs such that excess RPs bind to Mdm2, a p53 ubiquitinating protein. This interaction prevents Mdm2 binding to and ubiquitinating p53 for proteasomal degradation. Consequently, p53 accumulates resulting in apoptosis of neural crest cells, the precursors of most of the craniofacial skeleton^25^.

Homozygous null mutations in Pol I subunits, and other genes required for rRNA transcription leads to preimplantation lethality in mouse^25–28^. While the mechanism for cell death is most likely p53 dependent^26^, it is unknown if nucleolar structure is affected by the loss of Pol I function and rRNA transcription. Prior to fertilization, oocytes lack the classic tripartite nucleolar organization. However, following fertilization, at the 2-cell stage, when the zygotic genome is activated and rDNA transcription begins nucleolar precursor bodies (NPB) appear which have a compact fibrillar structure. At embryonic day (E) 2.5, when mouse embryos have 8-16 cells, NPBs begin to form small FCs that are partially surrounded by a DFC. The tripartite structure of the nucleoli is observed from the blastocyst stage at 3.5 dpc onwards (reviewed in ^29^). Considering that nucleologenesis is dependent upon active rRNA transcription, we hypothesized that nucleolar structure would be altered in the Pol I mutants and that this alteration would lead to nucleolar stress and embryonic lethality.

Here we show that Pol I mutants (*Polr1a*^*-/-*^, *Polr1b*^*-/-*^, *Polr1c*^*-/-*^ and *Polr1d*^*-/-*^) have structural defects in the nucleolus during preimplantation development. In all Pol I mutants, there is a significant decrease in number of NPBs and a concomitant increase in nucleolar volume. We observe that upon pharmacological inhibition of Pol I, using BMH-21, preimplantation embryos exhibit two distinct phenotypes: a single nucleolus similar to Pol I mutants or fragmented nucleoli like bodies. We also observe the same two phenotypes in human induced pluripotent stem cells (hiPSCs) fluorescently tagged for each nucleolar compartment. We determined that the single condensed nucleolus phenotype is likely caused by changes in the LLPS properties of NPM1 in the GC causing a single round nucleolus to form. Furthermore, the fragmented nucleolar phenotype is caused by failure of the nucleolus to reform following mitosis due to inhibition of Pol I activity. Overall, these results suggest that perturbation of Pol I function, rRNA transcription and protein distribution within the nucleolus can affect the LLPS of its components resulting in changes in nucleolar structure, all of which leads to embryo lethality or the pathogenesis of ribosomopathies.

## Results

### Loss of function of Pol I subunits results in nucleolar defects

Mutations in Pol I subunits result in preimplantation lethality^25–27^. To understand the function of Pol I subunits during mouse preimplantation development, we utilized our previously generated *Polr1a*^*-/-*^, *Polr1c*^*-/-*^ and *Polr1d*^*-/-*^ null mouse mutants^25^ together with a new *Polr1b*^*-/-*^ null mutant mouse, collectively referred to as Pol I mutants hereafter. While control embryos develop to the blastocyst stage, the Pol I mutant zygotes undergo four rounds of cell division following fertilization before arresting at the 16-cell stage (Fig. 1A). Pol I mutant embryos fragment and appear as a ball of cells with no distinct boundaries between individual blastomeres (Fig. 1B).

**Figure 1.**
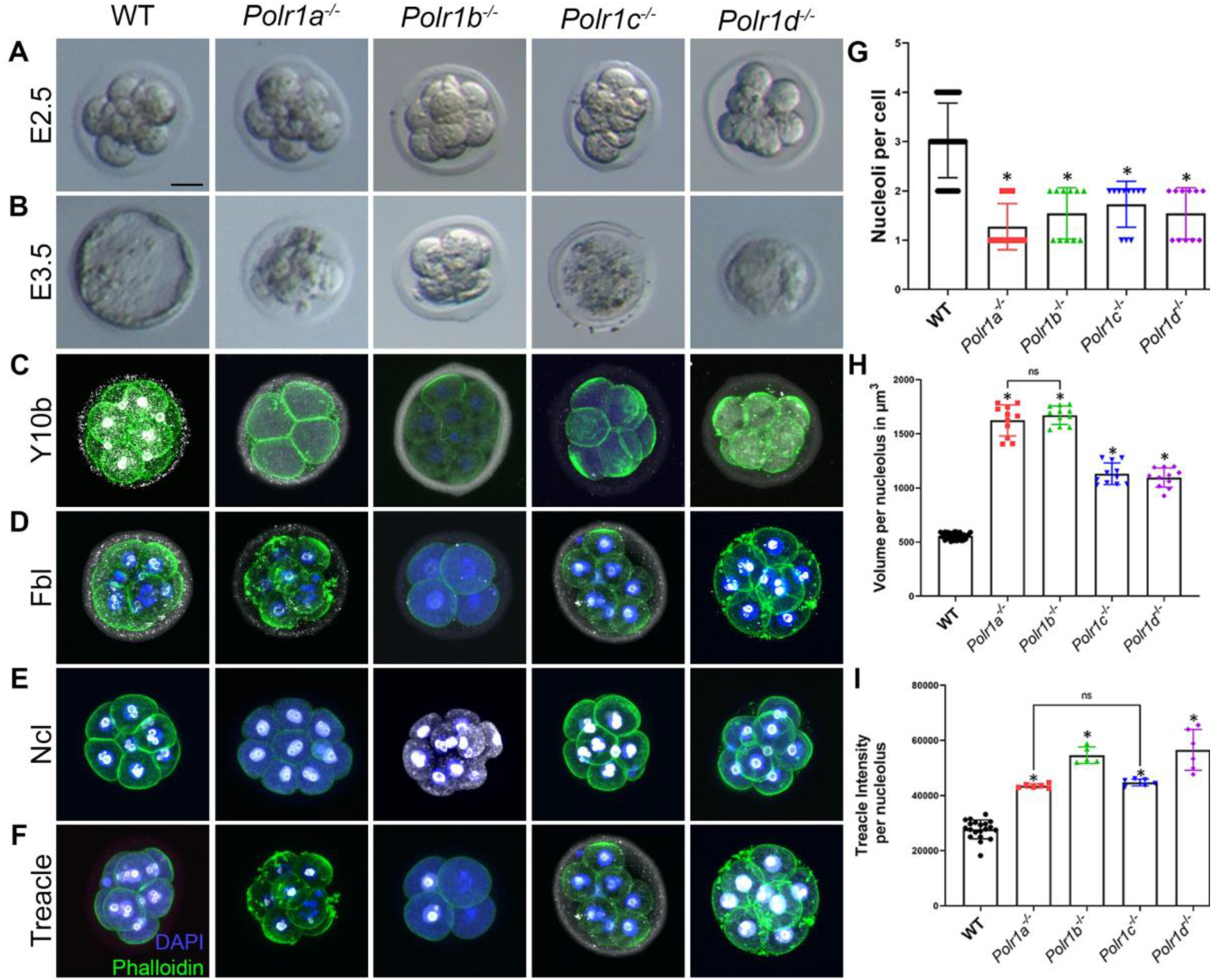
Loss of function of Pol I subunits results in nucleolar defects. Bright field images of WT, *Polr1a*^*-/-*^, *Polr1b*^*-/-*^, *Polr1c*^*-/-*^ and *Polr1d*^*-/-*^ embryos at E2.5 (A) indicate that the mutant embryos are indistinguishable from WT embryos at 8-16 cell stage. However, by E3.5 (B), Pol I mutant embryos arrest in association with blastomere fragmentation while the WT progresses to blastocyst stage. Immunostaining with Y10b to visualize rRNA (C) indicates that rRNA expression is reduced in all the mutant embryos. While Fbl (D) expression is variable, it is expressed in the NPBs of all the mutants. Ncl (E) and Treacle (F, I) expression is increased in the NPBs of all the mutants. In addition, the number of NPBs per blastomere is reduced in all the mutants (G), while the volume per NPB increases (H). The data is represented as mean+/-SEM. Scale bar for A-F = 12.5 μm.

rRNA transcription is required for nucleolar assembly. We therefore hypothesized that NPB structure would be disrupted in Pol I mutants. To assess for changes in NPB structure, we immunostained the Pol I mutants with markers which label rRNA and the three compartments of the nucleolus. We used the Y10b antibody to label 5.8S rRNA in the nucleolus as well as in the cytoplasm, and we observed a general downregulation of rRNA in Pol I mutant embryos (Fig. 1C). We used Fbl, Treacle and Ncl expression to demarcate the FC, DFC and GC, respectively. We observe that the level of Fbl expression is variably affected within the different NPBs (Fig. 1D), while Treacle and Ncl intensities are significantly and consistently increased within the NPB of Pol I mutants (Fig. 1 E, F, I). In addition, we find the number of NPBs per cell is decreased (Fig. 1G) and the volume of NPB is increased (Fig. 1H) in Pol I mutants compared to wildtype embryos. These phenotypes are indicative of perturbed nucleolar structure in the absence of Pol I activity.

### *Tcof1*^*-/-*^ mutants survive until mid-gestation

Tcof1/Treacle is a Pol I associated protein required for rRNA transcription^25,30^. Furthermore, mutations in *TCOF1* in humans result in Treacher Collins syndrome, similar to mutations in Pol I subunits^31–33^. We therefore hypothesized that *Tcof1* loss-of-function would also result in preimplantation lethality along with perturbed nucleolar structure^25^. Surprisingly, *Tcof1*^*-/-*^ embryos survive until E10.5, and although they have a heartbeat, they are grossly developmentally abnormal. At E7.5, *Tcof1*^*-/-*^ embryos are smaller compared to wildtype littermates (Fig. S1A). By E8.5, *Tcof1*^*-/-*^ embryos have a distinctly smaller head as well as delayed chorioallantoic fusion, which is indicative of defects in both ectoderm and mesoderm development (Fig. S1B). Given the survival of *Tcof1*^*-/-*^ embryos beyond gastrulation and the essential roles of Tcof1/Treacle in rRNA synthesis and DNA damage repair^25,34,35^, we hypothesized that Tcof1/Treacle may be maternally deposited and thus maternal protein activity continues in spite of the deletion of *Tcof1* in these embryos.

To test this hypothesis, we injected an antibody against Treacle into one blastomere of two-cell embryos to block the function of Treacle protein. These embryos survive until the late blastocyst stage, but in contrast to control embryos fail to hatch (Fig. S1C, D). We observe downregulation of Treacle and 5.8S rRNA (as measured by Y10b expression) as well as a single nucleolus phenotype in these Treacle knockdown embryos (Fig. S1E-H). In contrast to Pol I mutants, the expression of both Fbl and Ncl is significantly reduced in Treacle antibody injected blastomeres (Fig. S1I-L) and we observe a similar downregulation of nucleolar protein expression in *Tcof1*^*fx/fx*^*;Cre-ER*^*T2*^ mouse embryonic fibroblast cells (MEFs) (Fig. S2). Altogether this suggests that rRNA transcription is essential for nucleolar organization and structure.

### Nucleolar ultrastructure is altered in the *Polr1c*^*-/-*^ mutants

To better understand the effect of diminished Pol I function and rRNA transcription on nucleolar structure, we performed immunoelectron microscopy with Y10b and Ncl on *Polr1c*^*-/-*^ mutants. We observed that the number of FC and DFC were significantly reduced in individual blastomeres (Fig. 2A-A’). The staining patterns show that both rRNA and Ncl are redistributed to the edges of the GC, and that rRNA is diminished in the *Polr1c*^*-/-*^ mutants (Fig. 2B). Considering the FC and DFC of the NPBs are reduced in number and exhibit altered structure in *Polr1c*^*-/-*^mutants, we hypothesized this occurred in association with altered localization of active rDNA.

**Figure 2.**
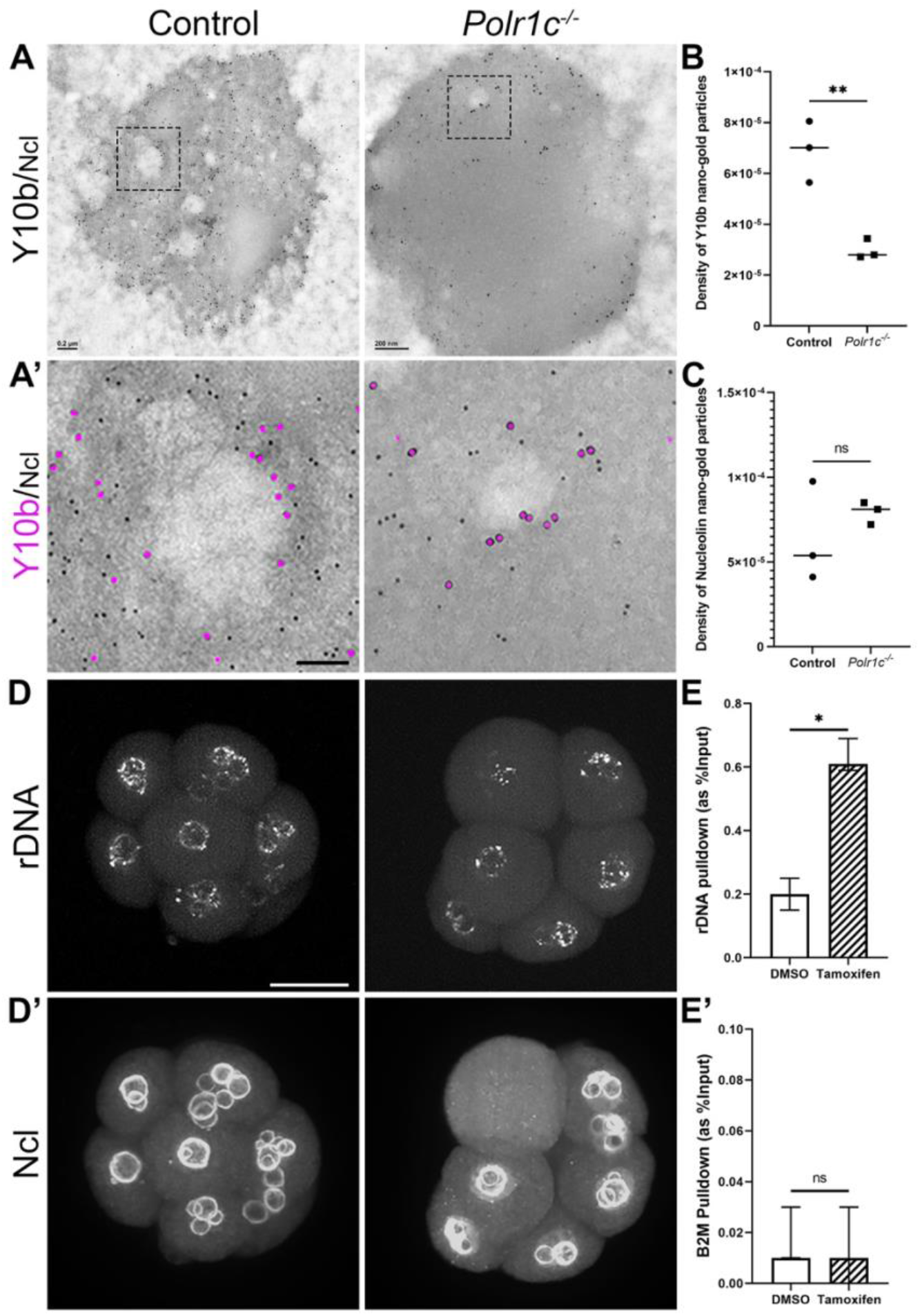
Nucleolar organization is altered in the *Polr1c*^*-/-*^ mutants. (A) Immunoelectron microscopy with antibodies against Y10b (large particles) and Ncl (small particles) on *Polr1c*^*-/-*^ mutant embryos indicate that the number of FC and DFC are significantly reduced in *Polr1c*^*-/-*^ mutants compared to control embryos. Scale bar = 200 nm. (A’) is higher magnification image of the FC/DFC shows lower levels of both Y10b (magenta) and Ncl (black) in the *Polr1c*^*-/-*^ mutants. Scale bar = 75 nm. Quantification (Mean+/-SEM) indicates that while Y10b is significantly reduced in *Polr1c*^*-/-*^ mutants (B), Ncl is not (C). 3D-immuno FISH with Ncl antibody (D’) and 5’ETS region of the 47S rDNA (D) indicates rDNA localization to the condensed nucleolus. Scale bar = 15 μm. (E) ChIP with Treacle antibody pulls down a significantly higher amount of rDNA promoter in tamoxifen treated *Polr1c*^*fx/fx*^*;Cre-ER*^*T2*^ MEFs compared to DMSO treated MEFs. B2M promoter was used as an internal control (E’). Data is represented as mean +/- upper and lower limits.

Fluorescence in situ hybridization with probes designed to the 5’ETS of rDNA followed by immunofluorescence for Ncl revealed that rDNA loci were redistributed within the nucleus around the single nucleolus in *Polr1c*^*-/-*^ mutants (Fig. 2D-D’), in a pattern reminiscent of nucleolar stress caps^36^.

As described above, the nucleolar disruption in Pol I mutants is accompanied by increased expression of Treacle in the NPB (Fig. 1F, I). We hypothesized that in the absence of Pol I on the rDNA promoter, more Treacle binds to the rDNA promoter either as a compensatory mechanism or as a means to prevent DNA damage^30^. However, the amount of biological material from preimplantation embryos is too low to perform a chromatin immunoprecipitation (ChIP) assay, therefore we used MEFs generated from *Polr1c*^*fx/fx*^*;Cre-ER*^*T2*^ mice. We first confirmed that the nucleolar defects in *Polr1a*^*fx/fx*^*;Cre-ER*^*T2*^ and *Polr1c*^*fx/fx*^*;Cre-ER*^*T2*^ MEFs are similar to *Polr1a*^*-/-*^ and *Polr1c*^*-/-*^ embryos (Fig. S2). Next we performed ChIP with an antibody against Treacle with lysates prepared from *Polr1c*^*fx/fx*^*;Cre-ER*^*T2*^ MEFs treated with DMSO or Tamoxifen. qPCR after ChIP revealed a significantly higher amount of 47S rDNA promoter was pulled down with Treacle in Tamoxifen treated *Polr1c*^*fx/fx*^*;Cre-ER*^*T2*^ MEFs compared to DMSO treated MEFs. Altogether, this suggests that in the absence of Pol I activity, the cells try to compensate with other proteins to maintain rRNA transcription, yet fail, leading to nucleolar disruption.

### Pharmacological inhibition of Pol I activity results in two phenotypes

Pol I mutants exhibit a consistent reduction in the number of NPB that coalesce to a single large nucleolus. To unveil the mechanisms underlying this phenotype we treated preimplantation embryos with BMH-21, a Pol I specific inhibitor^37,38^. Following treatment of one-or two-cell embryos with any concentration of DMSO or BMH-21, we observe embryo arrest at the 2-cell stage in association with blastomere fragmentation. Similarly, 4-cell stage embryos treated with BMH-21 fail to progress to the 8-cell stage. In contrast to control DMSO treated embryos which form blastocysts, 8-cell stage embryos treated with BMH-21 undergo one round of cell division prior to embryo lethality. When these BMH-21 treated 8-cell stage embryos reached the 16-cell stage, we observed two distinct phenotypes: a single nucleolus (Fig. 3B, yellow arrow) as was expected from our previous Pol I data; and a fragmented nucleolus (Fig. 3B, red arrow). The fragmented nucleolus manifests as speckles of Treacle, Ncl and Y10b throughout the nucleoplasm. Interestingly, E8.5 embryos treated with BMH-21 also present with a similar single nucleolus and fragmented nucleoli phenotypes (Fig. 3C-D). Altogether, these data suggest that Pol I inhibition results in two distinct nucleolar phenotypes and we next aimed to uncover the mechanism underlying these phenotypes.

**Figure 3.**
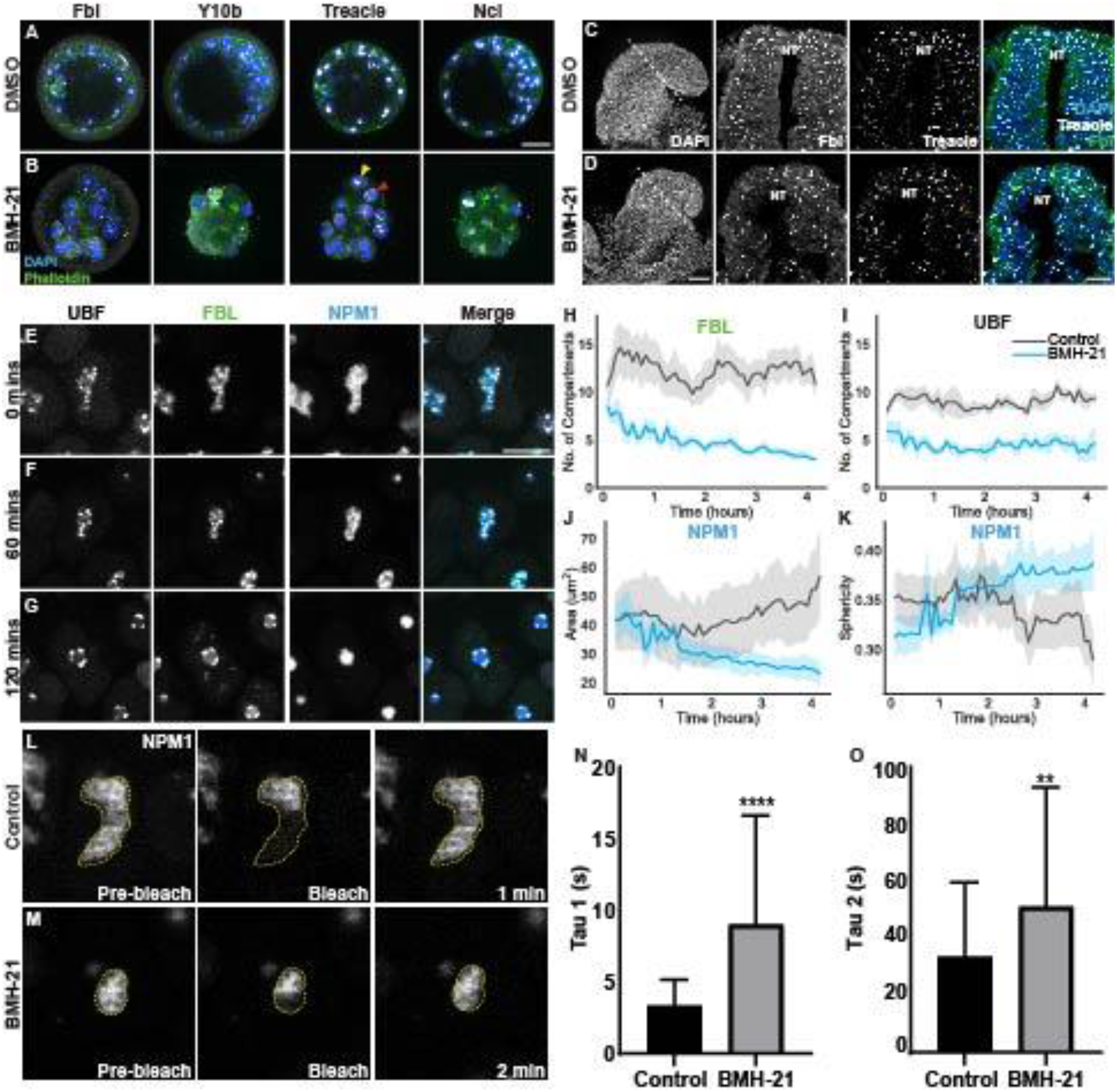
Pol I inhibition alters nucleolar structure and LLPS properties. (A, B) 16-cell stage DMSO and BMH-21 treated mouse embryos immunostained with Fbl, Y10b, Treacle and Ncl and counterstained with DAPI and Phalloidin. The yellow arrow indicates the condensed nucleolar phenotype, while the red arrow points to the fragmented nucleolar phenotype. Scale bar = 20μm. (C, D) E8.0 embryos treated with DMSO and BMH-21 immunostained with Fbl and Treacle, and counterstained with DAPI. Scale bar = 50μm. (E-G) Stills from live imaging video of fluorescently tagged hiPSCs treated with BMH-21 showing the nucleolus condensing. Scale bar = 10μm. (H-K) Analysis of live imaging videos showing changes in the number of UBF (H) and FBL (I) puncta, NPM1 area (J), and NPM1 sphericity (K). n =15 cells. Data is represented as mean (line) +/- 95% CI (shading). (L-M) Stills from half FRAP time-lapse video of pre-bleach, bleach, and recovery of NPM1 in control (L) and BMH-21 treated (M) cells. (N-O) Graphs of Tau values for half FRAP (N) and whole FRAP (O) showing both Tau1 and Tau2 significantly increase upon BMH-21 treatment. The data is represented as mean+/-SD.

### Pol I inhibition alters nucleolar structure and LLPS properties

The single nucleolus phenotype has previously been attributed to activation of the nucleolar stress pathway^39^, but how it occurs remains poorly understood. To better understand the real time dynamics and biophysical properties that underlie this phenotype, we performed live imaging of hiPSCs in which each compartment of the nucleolus is fluorescently tagged. Halo-tagged UBF labels the FC, GFP-tagged Fibrillarin (FBL) demarcates the DFC and an RFP-tagged Nucleophosmin1 (NPM1) marks the GC, allowing for real-time visualization of the dynamic morphology of each nucleolar compartment.

First, we tested if inhibiting Pol I transcription using BMH-21 results in a nucleolar phenotype similar to that observed in Pol I mutant preimplantation embryos. Using live imaging of BMH-21 treated cells, we observed a reduced number of FCs (UBF, grey) and DFCs (FBL, green) which instead formed characteristic nucleolar caps around the GC (NPM1, blue) which collectively coalesces into a single round nucleolus (Fig. 3E-I, Video 1)^39^. In the GC, the NPM1 area decreases, and sphericity increases following BMH-21 treatment (Fig 3G, J, K), similar to the coalesced single nucleolus we observed in BMH-21 treated preimplantation embryos (Fig. 3B, yellow arrow). These results indicate that when Pol I activity is perturbed hiPSCs present with a similar nucleolar phenotype as BMH-21 treated and Pol I mutant preimplantation embryos.

Nucleolar structure and Pol I function are intimately linked in that perturbations in one affect the other. We therefore hypothesized that the perturbation in nucleolar structure was caused by changes in the LLPS properties of the nucleolus upon loss of rRNA transcription. To test whether the LLPS properties of the nucleolus changed with Pol I inhibition, we performed half florescence recovery after photobleaching (FRAP) of NPM1 in the GC. This method is frequently used to measure changes in liquid-like droplets *in vivo*^40,41^. For half FRAP, half of the NPM1 area is photobleached and monitored for fluorescence recovery (Fig. 3L-M, Video 2). A fast half FRAP recovery would suggest liquid-like properties (i.e. low viscosity) while a slow half FRAP recovery indicates more gel-like properties (i.e. high viscosity)^40,41^. Additionally, full FRAP was performed to account for the addition of nucleoplasmic NPM1 to the nucleolus (Fig. S3 C-H). The half FRAP recovery (Tau1) of BMH-21 treated cells (average Tau1= 9.05s) is significantly longer than control cells (average Tau1= 3.33s), indicating slower NPM1 recovery in the GC. These results suggest the viscosity of the GC significantly increases with Pol I inhibition (Fig. 3N). The whole FRAP recovery (Tau2, Fig 3O) also increases with BMH-21 treatment, suggesting there is reduced affinity of NPM1 for the GC underlying the increased recovery. The whole FRAP percent recovery (Fig. S3B) decreases slightly with BMH-21 treatment, indicating there is less nucleoplasmic NPM1 available to add to the nucleolus. Overall, our data suggests the single nucleolar phenotype is likely a result of the changes in the LLPS properties of the nucleolus, specifically the GC, precipitated by the perturbation of Pol I function and decreased rRNA transcription.

### BMH-21 treatment prevents nucleolar reassembly following cell division

In addition to the coalesced single nucleolus phenotype, we also observe a fragmented nucleolar phenotype in a subset of cells in BMH-21 treated preimplantation embryos (Fig. 3B, red arrow), midgestation embryos (Fig. 3D) and hiPSCs (Fig. 4L). Live imaging of control hiPSCs indicates that during mitosis the nucleolus disassembles leaving UBF puncta associated with rDNA (Fig. 4A-C). Following telophase, the nucleolus begins reassembly with DFC and GC components, such as FBL and NPM1, first aggregated into prenucleolar bodies (PNBs) in the nucleoplasm (Fig. 4D). Once Pol I transcription restarts, the PNBs are recruited to the sites of Pol I transcription and the nucleolus reforms (Fig. 4E-F, Video 3). In cells treated with BMH-21, the nucleolus disassembles similar to control cells (Fig. 4G-I), however following mitosis the nucleolus is unable to reassemble (Fig. J-L, Video 3). This results in a fragmented nucleolar phenotype where individual UBF, FBL, and NPM1 puncta disperse throughout the nucleoplasm (Fig. 4L). To test if the fragmented phenotype is primarily observed in cells unable to reform their nucleolus following cell division when Pol I activity is inhibited, we synchronized cells in G2/M using VM-26^42,43^ and then released the cells with or without BMH-21 treatment. Following synchronization, cells were immediately fixed or released into regular media, we observed 0% of cells with a fragmented phenotype (Fig. 4N, O, Q). BMH-21 treatment after synchronization resulted in a significant increase in the percentage of cells (14.4%) with fragmented nucleoli (Fig. 4P-Q) compared to the 7.5% of cells with fragmented nucleoli in unsynchronized BMH-21 treated cells (Fig. 4M). These results suggest that inhibition of Pol I activity results a fragmented nucleolar phenotype following mitosis.

**Figure 4.**
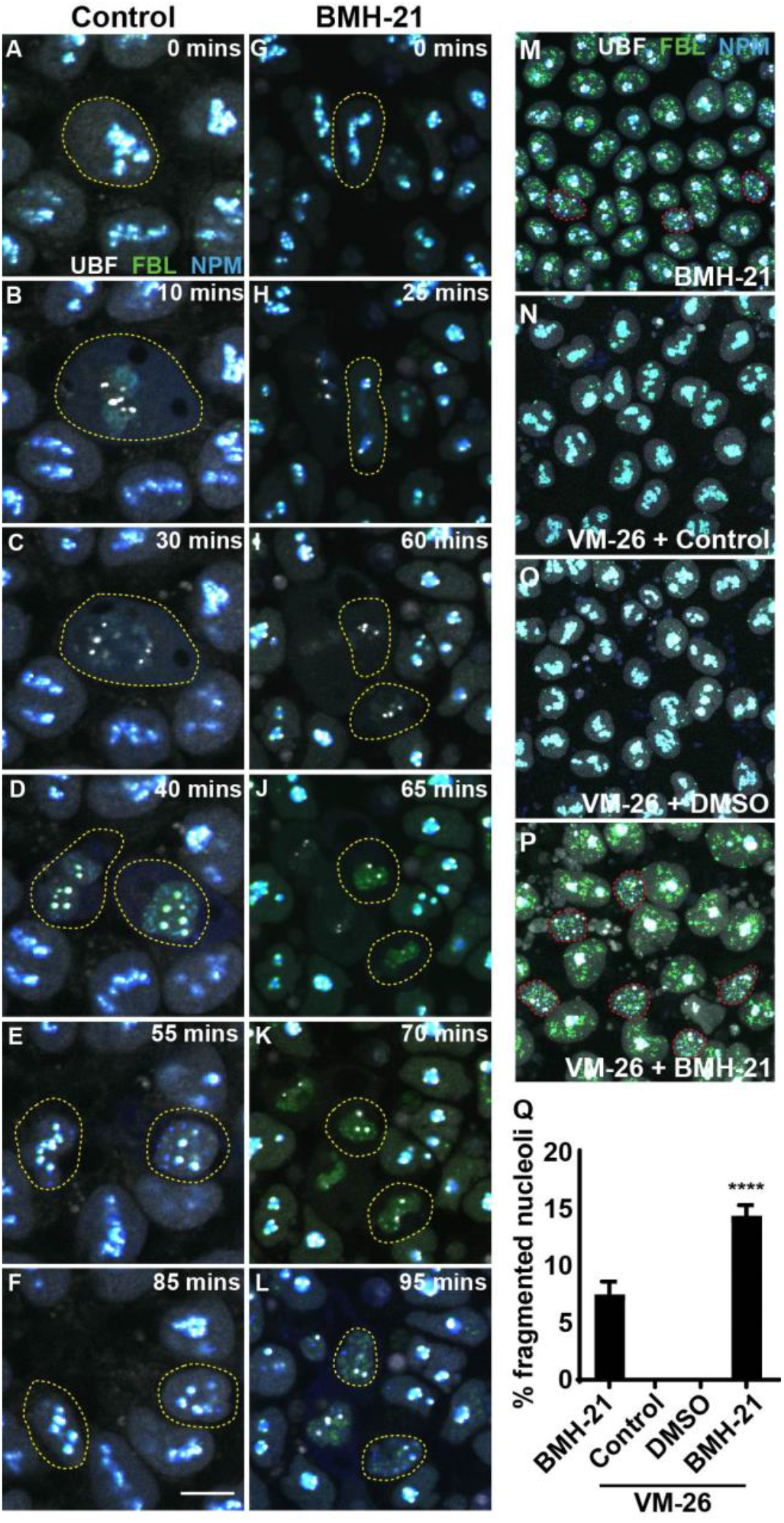
BMH-21 inhibition of Pol I function prevents nucleolar reassembly following cell division. (A-L) Stills from time-lapse imaging of a control (A-F) and BMH-21 treated (G-L) hiPSCs progressing through the cell cycle. UBF = grey. FBL = green. NPM1 = blue. Yellow lines denote dividing cells. Scale bar = 10μm. (M-P) Confocal images of hiPSCs treated with BMH-21 (M), synchronized in G2/M with VM-26 (N), synchronized with VM-26 and released for 3 hours in regular media (O), and synchronized with VM-26 and released for 3 hours with BMH-21 treated media (P). Red lines denote cells with fragmented nucleoli. (Q) Graph of the percentage of cells with fragmented nucleoli indicating that BMH-21 treatment just prior to cell division increases percentage of cell with fragmented nucleoli. Data is represented as mean+/-SD.

Taken together, our data indicate that Pol I inhibition affects nucleolar structure, however the specific phenotype depends on the cell cycle phase. If cells are in G0/G1/S, their nucleoli coalesce into a large single nucleolus. However, if cells are in G2/M phase, their nucleoli fragment, the difference being active rRNA transcription, which is essential to seed nucleolus reformation.

## Discussion

The nucleolus is the largest membrane-less organelle in an eukaryotic cell and it has a well-established role in ribosome biogenesis and several cellular stress responses, including DNA damage and proteotoxic stress^44^. Nucleolar dysfunction is linked to several human congenital diseases^1,20,23,45–48^, cancer^49,50^, neurodegenerative diseases^51,52^, viral infections^53–55^ and aging^56^.

The assembly of the nucleolus is dependent on actively transcribing rDNA^57,58^. Pol I inhibition in mouse zygotes using CX-5461 and BMH-21 has been previously shown to result in 2-cell or 4-cell stage embryo lethality^59,60^. However, Pol I mutant embryos, in which rRNA is genetically downregulated, survive until the 16-cell stage, leading us to question how these embryos survive an extra day and what happens to their nucleolar structure. Pol I mutant blastomeres present with a reduced number of NPBs and significantly increased volume of NPB. We hypothesize that these embryos survive until 16-cell stage despite the loss of rRNA transcription due to maternal deposits of rRNA, ribosomes^28,61^ and proteins similar to the Subcortical Maternal Complex^62–65^.

While the single nucleolus phenotype has previously been described as a consequence of perturbed ribosome biogenesis^66^, the dynamic changes in structure and their underlying biophysical reasons were unknown. Deletion of Pol I subunits disrupts the transcription of rDNA, reducing the rRNA content and altering the localization of nucleolar proteins such as Ncl and Treacle. Considering ribonucleoprotein particles maintain the LLPS properties of the nucleolus, we evaluated changes in the LLPS properties of the GC upon treatment with BMH-21 using FRAP. Our results indicate that perturbation of Pol I function with BMH-21 increases the viscosity of NPM1 in the GC. NPM1 typically exists as a pentamer^15^ and contains both intrinsically disordered regions (IDRs)^67,68^ that allow for homotypic interactions and an RNA binding domain (RBD) which binds to the processed rRNA entering the GC^15,68,69^. Under normal conditions, NPM1 pentamers bind to rRNA through its RBD and assist in pre-ribosomal assembly until the ribosomal subunits are expelled from the GC^70^. However, when Pol I transcription is disrupted, which reduces the amount of rRNA entering the GC, this decreases NPM1-rRNA interactions and increases NPM1 homotypic interactions^70,71^. The increase in NPM1 homotypic interactions increases the density of the NPM1 meshwork and consequently GC viscosity^6,67,69^, resulting in a single nucleolus. Additionally, the single large nucleolus phenotype is consistent with the observation that the number of nuclear speckles, another membrane-less organelle in the nucleus, decreases upon transcriptional inhibition^72^. Therefore, inhibiting rDNA transcription leads to changes in NPM1 interactions in the GC and subsequently altered nucleolar structure^71^.

While nucleolar proteins, such as NPM1, can drive phase separation *in vitro* independently^6^ rRNA also aids in maintaining LLPS of the nucleolus^7374^ and can alter LLPS droplet viscosity *in vitro*^7574^ and promote the immiscibility of the DFC and GC through interaction with FBL and NPM1^6,15,73^. Our data suggests that reducing rRNA by inhibiting Pol I limits NPM1 interactions with rRNA and other NPM1 molecules leading to redistribution of NPM1 and altered LLPS properties of the GC (Fig. S4). Interestingly, we also observe drastic relocalization of Ncl and Treacle in our Pol I mutant preimplantation embryos. Ncl contains a Gly-Arg-rich (GAR) domain (similar to FBL’s IDR^76,77^) and Treacle is also an intrinsically disordered protein^78^. Therefore, their change in localization could be due to changes in their interactions with rRNA or other proteins in the DFC, leading to changes in LLPS properties. Overall, these data demonstrate that impairing Pol I function and rRNA transcription leads to changes in the LLPS properties of the nucleolus, and this in turn alters nucleolar structure.

In both the hiPSCs and preimplantation embryos treated with BMH-21, we observed two distinct phenotypes, a single condensed nucleolus, and fragmented nucleoli. The fragmented phenotype results from the inability of the nucleolus to reform following cell division. Typically, the nucleolus disassembles at the beginning of mitosis upon cessation of Pol I transcription^79,80^ and then reassembles following the re-initiation of transcription^81^. Following cell division, UBF and pre-rRNA processing factors, such as NCL, are recruited to active NORs in early telophase^57,58^. Also, during early telophase, DFC and GC factors, such as FBL and NPM1, are localized in discreet puncta in the nucleoplasm called pre-nucleolar bodies (PNBs)^57,58^. Upon Pol I activation in late telophase, the PNBs are recruited to sites of active rRNA transcription and the nucleolus reforms. In BMH-21 treated hiPSCs, the nucleolus successfully disassembles preceding cell division, however, following cell division the nucleolus fails to reform and instead we observe discreet small puncta of UBF, FBL and NPM1 dispersed throughout the nucleoplasm. Our data demonstrates that when cells are synchronized in G2/M and Pol I activity is inhibited, an increased percentage of cells exhibit a fragmented nucleolar phenotype after division. This data suggests that the nucleolar phenotype, single condensed or fragmented, is dependent on the phase of the cell cycle when Pol I is inhibited.

The change in nucleolar structure in the absence of rRNA transcription most likely leads to nucleolar stress which is associated with p53 activation^25,26^. p53 activation in Pol I mutant and zebrafish embryos results in lethality^20,25–27,82^ and in human patients leads to ribosomopathies. While DNA damage has been suggested to occur following perturbation of Pol I activity, we find no evidence for this *in vivo* in our genetic or other analyses^25^. Furthermore, BMH-21 has been shown to elicit its effect on Pol I function, independent of any DNA damage response^37^. Overall, our work has uncovered the molecular and biophysical mechanisms underlying structural changes in the nucleolus resulting from genetic and chemical inhibition of Pol I function and rRNA transcription.

## Methods

### Animals

All animal experiments were conducted in accordance with Stowers Institute for Medical Research Institutional Animal Care and Use Committee approved protocol (IACUC #2022-014) and Virginia Commonwealth University Transgenic/Knockout Mouse Facility (Virginia Commonwealth University IACUC no. AM10025). The day a vaginal plug was observed in a time mated female was designated as embryonic day (E) 0.5. All mice were housed in a 16 hour light: 8 hour dark light cycle. *Polr1a*^*+/-*^, *Polr1c*^*+/-*^ and *Polr1d*^*+/-*^ mice were generated and maintained as previously described^25^. To generate *Tcof1*^*Δ/Δ*^, *Tcof1*^*fx/fx*^ female mice^25^ were crossed with Zp3-Cre line to delete *Tcof1* in germline cells.

CRISPR-Cas9 technology was used to engineer a new mouse strain containing a deletion of exon 2 and 3 in the *Polr1b* gene. Potential guideRNA target sites were designed using CCTOP^83^. The potential target sites were evaluated using the predicted on-target efficiency score and the off-target potential^84^. GuideRNA target sites were designed in intron 1 and intron 3 to fully delete exons 2 and 3. For both selected guideRNA targets, the sequence was ordered as Alt-R CRISPR-Cas9 crRNA from Integrated DNA Technologies (IDT). Each crRNA was hybridized with a universal tracrRNA (IDT) to form a full length guideRNA. Ribonucleoprotein (RNP) was prepared for microinjection with 10ng/μl of each full length guideRNA and 10ng/μl Cas9 protein (IDT, #1081059).

Tissue samples (ear and tail clips) from resulting animals were lysed using QuickExtract DNA Extraction Solution (Epicentre) followed by PCR at the specific genomic location. Amplification products were analyzed for a size shift using a LabChip GX (Perkin Elmer). Selected samples with deletion sized products were purified using ExcelaPure 96-Well UF PCR Purification Kit (Edge Bio) followed by Sanger sequencing.

### Brightfield imaging

Embryos were harvest at E2.5 by flushing the oviduct with 1ml M2 media as described previously^85^. The embryos were imaged using a Leica MZ16 microscope equipped with a Nikon DSRi1 camera and NIS Elements BR 3.2 imaging software.

### Immunostaining

The harvested embryos were transferred to a 4 well dish containing 4% PFA (Alfa Aesar via VWR Cat. No. AA43368-9M) for 10 mins. Immunostaining was performed as described previously^86^. The primary antibodies used are Tcof1 (Abcam, #ab65212), Y10b (Abcam, # ab171119), Nucleolin (Abcam, #ab22758) and Fibrillarin (Abcam, # ab154806). The volume and intensity of staining was measured using IMARIS. All images for each stain individually were acquired with the same settings and brightness and contrast adjusted the same throughout.

### Mouse embryonic fibroblast cells

Mouse embryonic fibroblasts were derived from E13.5 *Polr1a*^*fx/fx*^*;Cre-ER*^*T2*^, *Polr1c*^*fx/fx*^*;Cre-ER*^*T2*^ and *Tcof1*^*fx/fx*^*;Cre-ER*^*T2*^ embryos and cultured as described previously ^25^. Control cells were treated with DMSO, while mutant cells were treated with 5μM Tamoxifen. All experiments were performed 48 hours post tamoxifen treatment. All images for each stain individually were acquired with the same settings and brightness and contrast adjusted the same throughout.

### Immunoelectron microscopy

For immunogold labelling, embryos were fixed in 4% PFA, dehydrated in ethanol, and embedded in LR-White resin. Ultrathin sections of about 80nm in thickness were mounted on Formvar and carbon coated copper grids, then washed three times with PBS and three times with PBS containing 1% bovine serum albumin and 0.15% glycine, followed by 30 min blocking with 5% normal goat serum. Samples were incubated 1 hour with the Ncl and Y10b primary antibodies at room temperature. After washing in PBS, samples were incubated for 1 hour with gold-conjugated secondary antibodies (Jackson ImmunoResearch, #115-205-166 and 111-195-144). Sections were stained with 2% uranyl acetate and observed under a FEI electron microscope at 80 kV. The specificity of the immunoreaction was assessed in all cases by omitting the primary antibodies from the labelling protocol and incubating the sections only in the gold-conjugated secondary antibodies.

Quantification was performed as follows. To localize individual gold particles, Laplacian of Gaussian filters of an appropriate size, depending on the image resolution and particle size, were applied and maxima found using plugins in Fiji^87^. First the larger 12nm particles were found, and then regions around them were eliminated to prevent them from being found when searching a second time for 6nm particles. These were then masked against a manually annotated nucleolar outline and the particles/nm2 were computed for each image and particle size.

### 3D-immuno FISH

Embryos were fixed with 4% paraformaldehyde (PFA) for 10 min, rinsed and permeabilized with 0.5% TritonX-100/PBS for 60 min. RNA was digested with 200μg/ml of RNAse in PBS for 2 hours at 37°C, and embryos were rinsed with PBS two times. Blocking was performed using superblock (Thermo scientific, # 37580) for 2 hour at room temperature, and embryos were incubated with anti-Nucleolin antibody (Abcam, #ab22758) for 20 hours at 4°C. After three rinses with PBS, secondary antibody (Donkey Alexa 647 anti-Rabbit) (Invitrogen, #A-31573) was applied for 2 hours at room temperature and washed with PBS two times. Post fixation was performed with 4% PFA for 10 min, washed two times with PBS, and embryos were dehydrated with ice cold 100 % Methanol for 30 min. With small volume of 100% Methanol, embryos were deposited on microscope slides, and the slides were air dried for 1 hour. Probe DNA (mouse rDNA BAC clone RP23-225M6, Empire genomics) was mixed with Hybridization buffer (50% formamide, 2X SSC, 1% dextran sulfate, 100ug/ml salmon sperm DNA), applied on slides, and both genomic DNA and probe DNA were denatured simultaneously on a heating block at 80°C for 8 min. Hybridization was performed at 37°C for 16 to 20 hours. Slides were washed with 2X SSC for 5 min, 50% formamide/2X SSC for 15 min at 37°C, 2X SSC for 10 min two times, and 2X SSC/0.1% Triton X for 10 min. Embryos were stained with 10 ug/ml of DAPI for 30 min at room temperature. Slides were mounted with ProLong gold antifade mountant (Invitrogen, # 36930). The embryos were imaged using Nikon Ti2 with CSU-W1 Spinning Disk. All images for each stain individually were acquired with the same settings and brightness and contrast adjusted the same throughout.

### Chromatin Immunoprecipitation

To pulldown rDNA promoter, chromatin immunoprecipitation was performed as described previously^88^ using Treacle antibody. Student’s t-test was used for statistical analysis. Primers used are as follows: rDNA_F: 5’-TCTGGTACCTTCTTAATCACAGAT-3’; 5’-rDNA_R: ATAAATGAAGAAAATAACTAA-3’; B2M_F: 5’-CACTGACCGGCCTGTATGC-3’; B2M_R: 5’-CACTGACCGGCCTGTATGC-3’.

### BMH-21 treatment on preimplantation embryos

8-cell embryos from CD1 pregnant dam were harvested using M2 media and cultured in KSOM media in 5% CO_2_ at 37°C. After 30 minutes of normalization in the culture conditions, the embryos were treated with 0.1 μM BMH-21 for 8 hours. The controls were treated with DMSO. After 8 hours, the culture media was changed to fresh KSOM media and cultured for 16 hours. The embryos were then fixed and immunostained.

### BMH-21 treatment on E8.5 embryos

E8.5 embryos from C57Bl/6 pregnant dam were harvested and cultured in 50% rat seum in DMEM-F12 media in 20% CO_2_ at 37C. After 30 minutes of normalization in the culture conditions, the embryos were treated with 1 μM BMH-21 for 8 hours. The controls were treated with DMSO. After 8 hours, the embryos were then fixed and immunostained. Further, the embryos are cryo-sectioned and imaged using Nikon Ti2 with CSU-W1 Spinning Disk. All images for each stain individually were acquired with the same settings and brightness and contrast adjusted the same throughout.

### Antibody microinjections in 2-cell embryos

Immature C57BL/6J female mice (3-4 weeks of age) were utilized as embryo donors. The C57BL/6J females were superovulated following standard procedures with 5 IU PMSG (Genway Biotech, #GWB-2AE30A) followed 46 hours later with 5 IU hCG (Sigma, #CG5) and subsequently mated to fertile C57BL/6J stud males. Females were checked for the presence of a copulatory plug the following morning as an indication of successful mating. One-cell fertilized embryos at were collected from the oviducts of successfully mated females at 0.5dpc and placed in KSOM media in a CO_2_ incubator at 37°C, 5% CO2. Fertilized oocytes were cultured over-night to the two-cell stage for microinjection the following morning. Antibody reagents were microinjected (1-2 pico liters total of 1mg/ml Treacle antibody) into one blastomere of the developing two-cell embryo using previously described techniques ^85^. In brief, microinjection was performed using a Nikon Eclipse Ti inverted microscope equipped with Eppendorf TansferMan^â^ micromanipulators, Eppendorf CellTram Air^â^ for holding of embryos, and Eppendorf FemtoJet^â^ auto-injector. A small drop of M2 media was placed on a siliconized depression slide and approximately 20 -30 C57BL/6J oocytes were transferred to the slide for microinjection. The slide was placed on the stage of the microscope and oocytes were injected at 200x. Immediately following microinjection, the embryos were returned to the CO2 incubator in KSOM culture media and observed daily for embryo development. At blastocyst stage, the embryos were fixed and immunostained.

### Cell culture practices

Human induced pluripotent stem cell line used in this study, AICS-0086 cl.147, was an edited line generated as described previously^89,90^. This cell line can be obtained through the Allen Cell Collection (www.allencell.org/cell-catalog). Undifferentiated hiPSCs were maintained and passed on plates coated with hESC-qualified Matrigel (Corning #354277) in mTeSR1 (Stem Cell Technologies #85850) supplemented with 1% penicillin/streptomycin (ThermoFisher, #15070063). Cells were passaged approximately every 3-5 days (70-85% confluency) using Accutase (Gibco, #11105-01) to detach cells. Cells were plated in mTeSR1 + 1% P/S and 10 µM Rock Inhibitor (Y-27632, StemCell Technologies, #72308). A full detailed protocol for cell line maintenance can be found at www.allencell.org/sops SOP: WTC culture v1.7.pdf. For BMH-21 treatment, BMH-21 (Sigma, #SML1183) was added to mTeSR1 media at 1 µM for either 1 hour for fixed imaging or after the first acquired frame for live imaging. For VM-26 treatment, 80nM of VM-26 (Sigma, #0609) was added to media for 24 hrs to synchronize cells in G2/M before being rinsed off.

### Live imaging acquisition

Human induced pluripotent stem cells with RFP-tagged Nucleophosmin, GFP-tagged Fibrillarin, and Halo-tagged upstream binding factor (AICS-0086 cl.147) were plated on Matrigel coated Ibidi 35 mm µ-Dishes (Ibidi, # 81156). Before imaging, cells were treated with 200 nM of Janelia Fluor® HaloTag® Ligand 646 for 30 mins. The cells were then imaged using a CSU-W1 spinning disc (Yokogawa) coupled to a Ti2 microscope (Nikon) through a 60x Plan Apochromat objective (NA 1.45). Excitation of GFP occurred at 488 nm, RFP occurred at 561 nm, and Halo JF 646 occurred at 640 nm. The emissions were collected for 20 to 200 ms per frame through a standard filter onto a Flash 4 camera (Hamamatsu). Cells were imaged every 10 mins for 5 hrs. Great care was taken to reduce illumination as much as possible to avoid phototoxicity.

### Live imaging analysis

Individual cells were cropped from time-lapse videos and imported into Imaris (Bitplane, Inc.). A total of 12 untreated cells from 3 different experiments and 12 drug treated cells from 3 different experiments were analyzed. Surfaces were created for each component of the nucleolus using fluorescent labels and tracked over time. The number of FBL and UBF components, and the area and sphericity of the NPM1 component were averaged and plotted with the 95% confidence interval per time point for each treatment.

### FRAP acquisition and analysis

Human induced pluripotent stem cells with RFP-tagged Nucleophosmin, GFP-tagged Fibrillarin, and Halo-tagged upstream binding factor (AICS-0086 cl.147) were plated on Matrigel coated Ibidi 35 mm µ-Dishes (Ibidi, #81156). The cells were then imaged using a CSU-W1 spinning disc (Yokogawa) coupled to a Ti2 microscope (Nikon) through a 60x Plan Apochromat objective (NA 1.45). Excitation of RFP occurred at 561 nm, and the emission was collected for 50 to 200 ms per frame through a standard filter onto a Flash 4 camera (Hamamatsu). Photobleaching was achieved using a diffraction-limited 561 nm laser beam focused on the region of interest and scanned across that ROI until fluorescence was eliminated. The settings for bleaching were adjusted so that the bleaching step was nearly instantaneous (< 2 s). Multiple prebleach images of cells were acquired for 0.5 s without delay, and the recovery after bleaching was recorded every 1 s for 2 minutes. In separate instances, either the entire punctum was bleached (full FRAP) or only a part was bleached (half FRAP). Recovery curves and analysis were performed using in-house written plugins in Fiji (https://imagej.net/Fiji). First, the images were cropped to the cell of interest and then registered to remove the cell/punctum movement using a plugin called Stackregj. After this, an ROI was placed over the bleached portion of the cell and the mean intensity of the ROI was plotted using a plugin called “create spectrum jru v1.” Once all the curves for a particular condition were collected, the curves were all combined into one window using “combine all trajectories jru v1.” The curves were then all normalized to the min and max of each curve using “normalize trajectories jru v1.” The curves were then manually aligned in time so that the bleach point all aligned at the same timepoint. Finally, each curve was fit individually using “batch FRAP fit jru v1.” The fit parameters were then averaged to give the tau and percent recovery.

## Acknowledgments

The authors thank members of the Trainor lab for their insights and discussions. This work was funded by the Stowers Institute for Medical Research (P.A.T), American Association for Anatomy Post-Doctoral Fellowship (S.D.) and K99 (DE030972) from the National Institute for Dental and Craniofacial Research (S.D.). Original data underlying this manuscript can be accessed from the Stowers Original Data Repository at http://www.stowers.org/research/publications/LIBPB-1752.

## Supplementary Information

**Figure S1.**
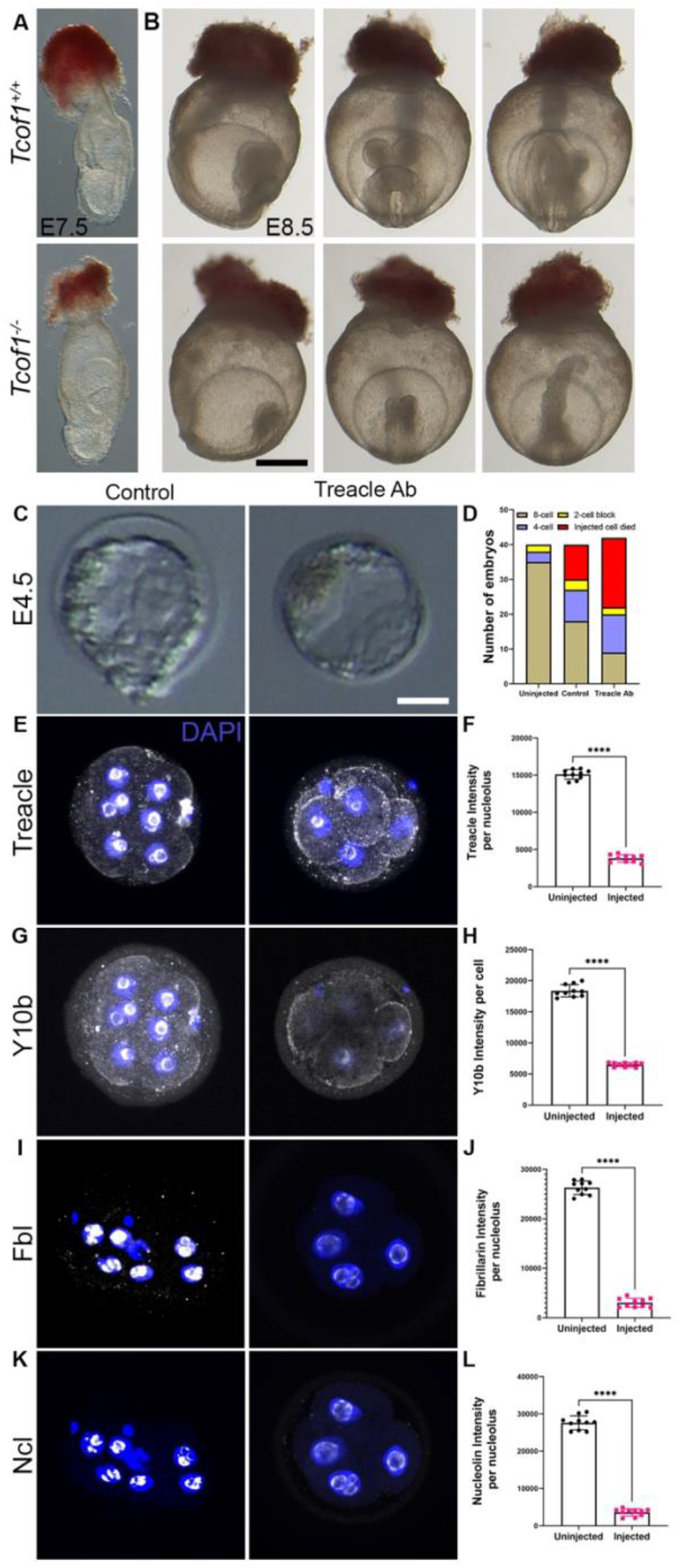
*Tcof1*^*-/-*^ embryos are midgestation lethal. (A) *Tcof1*^*-/-*^ have smaller embryo proper at E7.5. At E8.5 (B), these embryos are significantly smaller than the WT littermates. (C) Treacle antibody injected embryos have cell death in the inner cell mass, while control embryos proceed to hatching at E4.5 (D) Survival statistics of embryos injected with antibody indicates that over 50% of the embryos injected with Treacle antibody did not survive. Expression of Treacle (E, F), Y10b (G, H), Fbl (I, J) and Ncl (K, L) are significantly reduced in Treacle injected blastomeres and their derivatives. The data is represented as mean+/-SEM. Scale bar for A and B is 100 μm. Scale bar for E, G, I and K = 12.5 μm.

**Figure S2.**
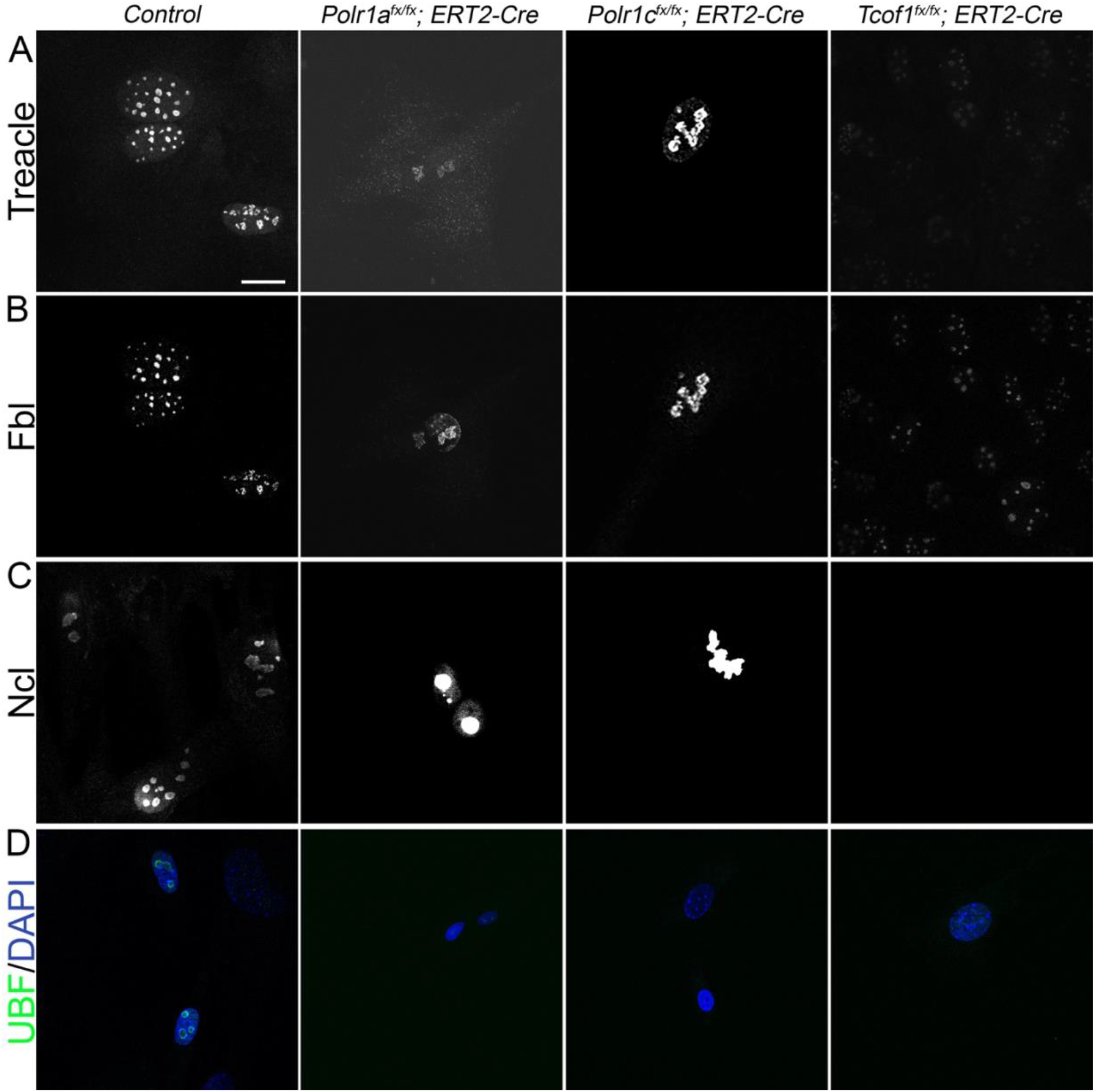
Mutations in Pol I subunits causes Nucleolar Morphology defects in MEFs. Immunostaining of MEFs with Treacle (A), Fbl (B), Ncl (C) and UBF (D) antibody suggests that the number of nucleoli significantly reduce in *Polr1a*^*fx/fx*^*;Cre-ER*^*T2*^, *Polr1c*^*fx/fx*^*;Cre-ER*^*T2*^ and *Tcof1*^*fx/fx*^*;Cre-ER*^*T2*^. *Tcof1*^*fx/fx*^*;Cre-ER*^*T2*^ MEFs have reduced expression of Fbl, Ncl and Ubf unlike *Polr1a*^*fx/fx*^*;Cre-ER*^*T2*^ and *Polr1c*^*fx/fx*^*;Cre-ER*^*T2*^.

**Figure S3.**
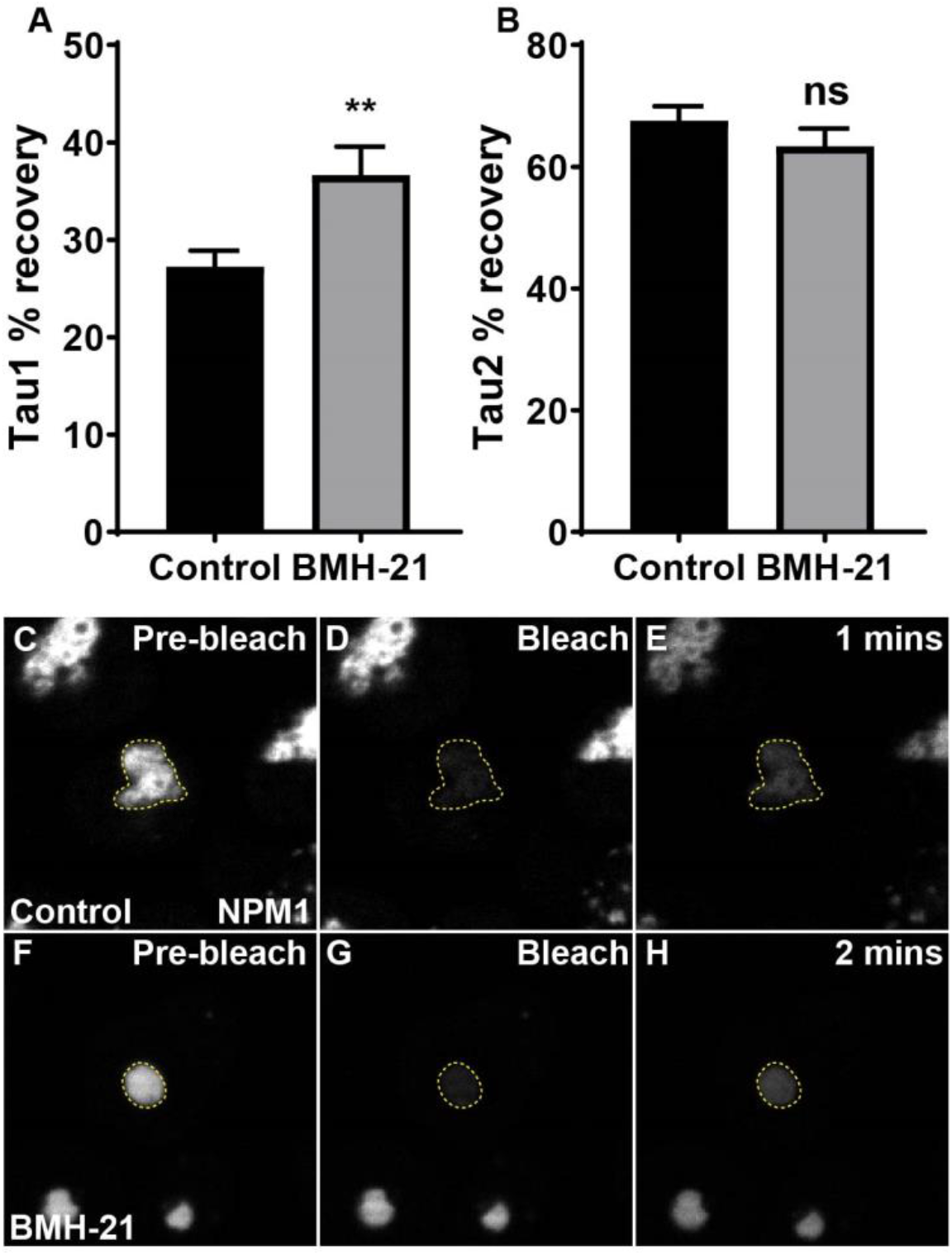
BMH-21 treatment causes increased recovery in whole and half FRAP GC. (A, B) Graphs showing the percent recovery of the GC of half and whole FRAP respectively. Data is represented as mean+/-SD. (C-H) Confocal images of whole FRAP NPM1 and their recovery over time in control and BMH-21 treated cells.

**Figure S4.**
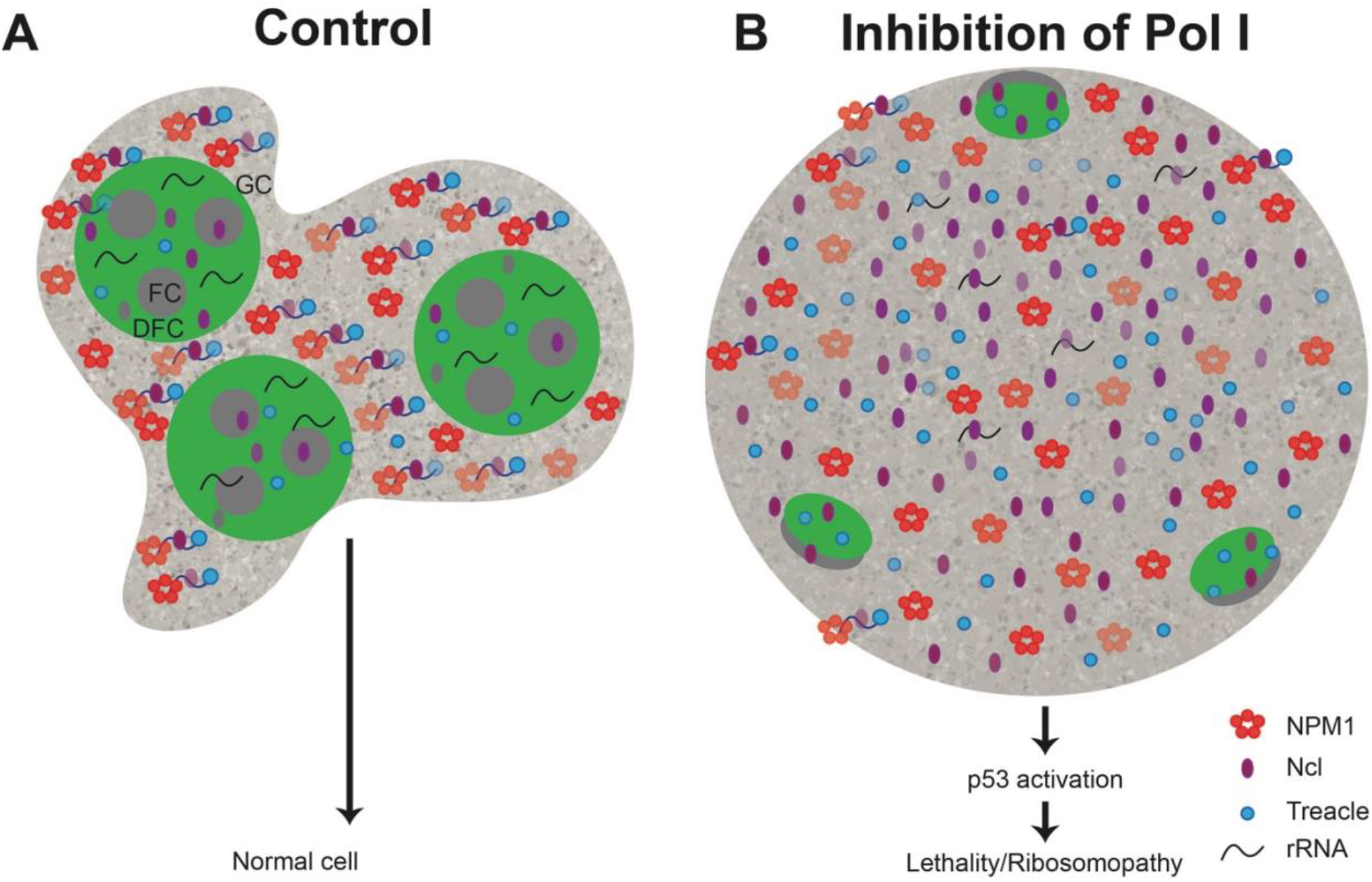
Nucleolar disruption in Pol I mutants. (A) In control embryos and cells, each nucleolus has distinct regions of FC and DFC surrounded by the GC and maintains an amorphous structure. rRNA is bound to nucleolar proteins (NPM1 in red, Ncl in purple and Treacle in blue). (B) When Pol I activity is inhibited, the nucleolus changes shape to round and condensed. rRNA transcripts are reduced and nucleolar protein expression increases, which leads to a change in the LLPS of the nucleolus. This leads to nucleolar stress and lethality/ ribosomopathy.

**Video 1.**
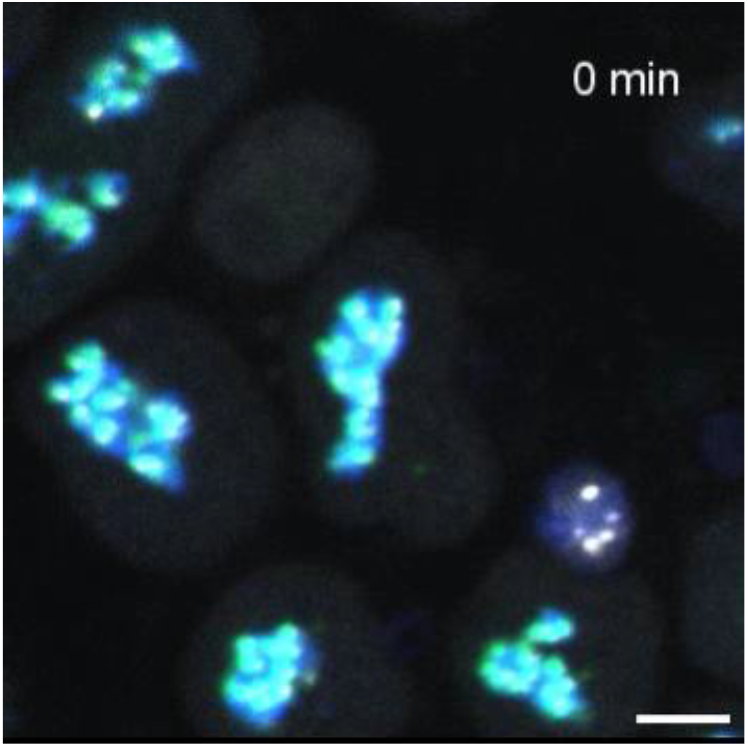
BMH-21 treatment of nucleolar tagged hiPSCs. hiPSCs were imaged for UBF (HaloTag grey), FBL (GFP, green) and NPM1 (RFP, blue) every 10 mins for 5 hours. 1μM of BMH-21 was added after 1^st^ frame acquired. NPM1 condenses into round circle while UBF and FBL form nucleolar caps at the GC periphery. Scale bar = 10μm.

**Video 2.**
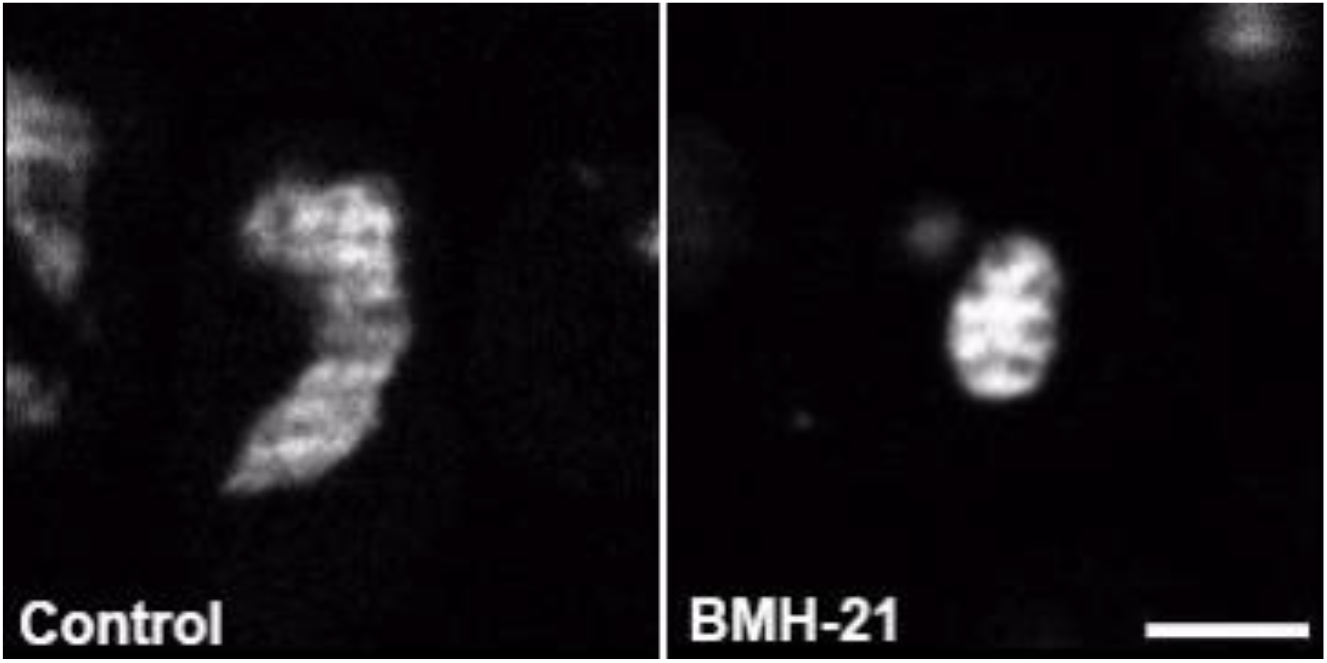
Half FRAP of control and BMH-21 treated hiPSC. NPM1 (RFP, grey) was imaged in control (left) and BMH-21 treated (right) rapidly for five frames pre-bleach, then half of the GC was bleached, and recovery of NPM1 was monitored for two minutes.

**Video 3.**
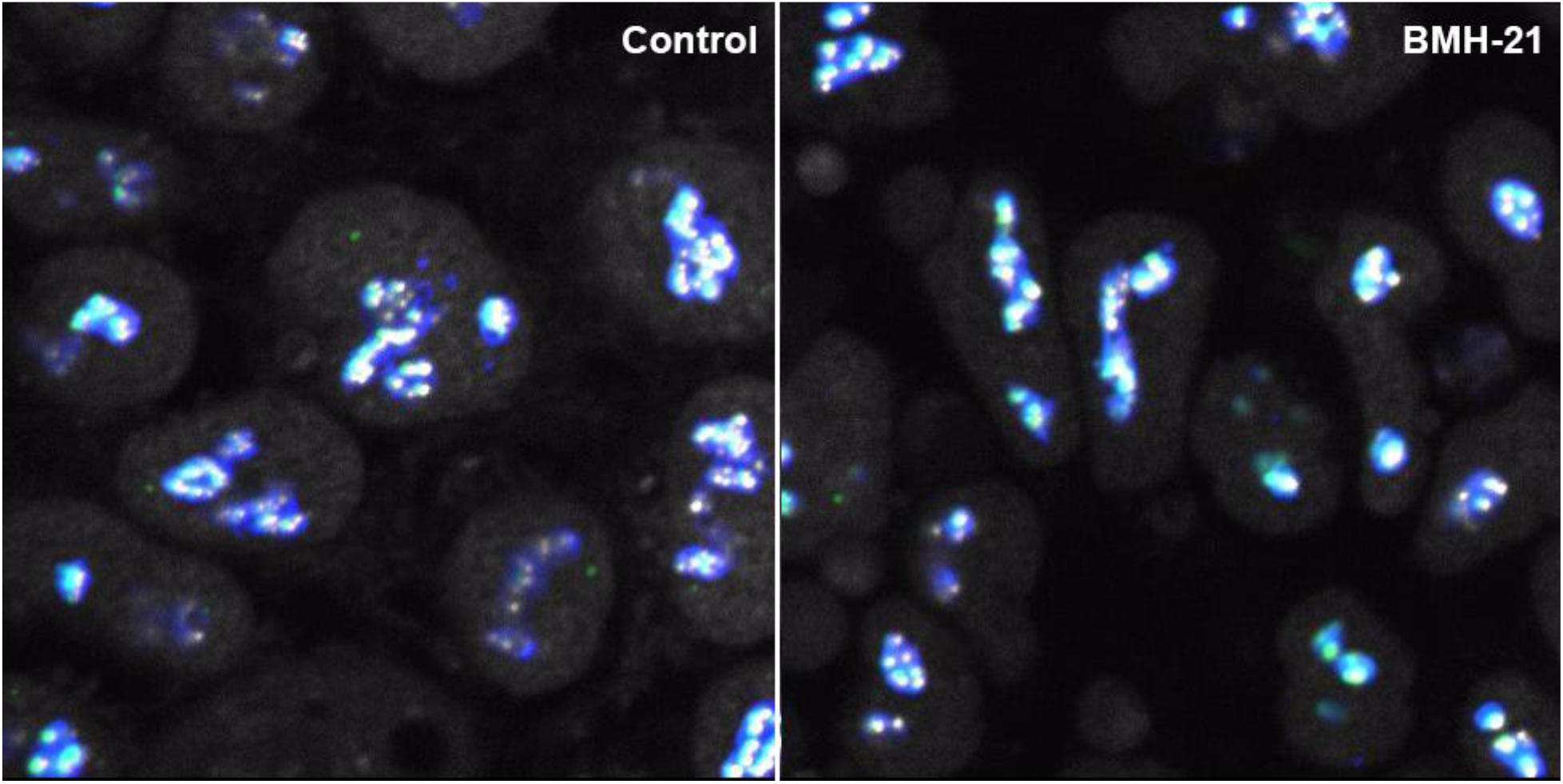
Cell division in control and BMH-21 treated hiPSCs. Time-lapse imaging of control (left) and BMH-21 treated (right) cells undergoing cell division. The nucleolus disassembles leading up to division with UBF remaining associated with the rDNA. In control cells, the nucleolus reassembles following division while the nucleolus in BMH-21 treated cell’s fragment.

